# Quantified Dynamics-Property Relationships: Data-Efficient Protein Engineering with Machine Learning of Protein Dynamics

**DOI:** 10.1101/2025.04.23.650227

**Authors:** T. Emme Burgin

## Abstract

Machine learning has proven to be very powerful for predicting mutation effects in proteins, but the simplest approaches require a substantial amount of training data. Because experiments to collect training data are often expensive, time-consuming, and/or otherwise limited, alternatives that make good use of small amounts of data to guide protein engineering are of high potential value. One potential alternative to large-scale benchtop experiments for collecting training data is high-throughput molecular dynamics simulation; however, to date this source of data has been largely absent from the literature. Here, I introduce a new method for selecting desirable protein variants based on quantified relationships between a small number of experimentally determined labels and descriptors of their dynamic properties. These descriptors are provided by deep neural networks trained on data from molecular dynamics simulations of variants of the protein of interest. I demonstrate that this approach can obtain very highly optimized variants based on small amounts of experimental data, outperforming alternative supervised approaches to machine learning-guided directed evolution with the same amount of experimental data. Furthermore, I show that quantified dynamics-property relationships based on only a handful of experimentally labeled example sequences can be used to accurately predict the key residues that are most relevant to determining the property in question, even when that information could not have been known or predicted based on either the molecular dynamics simulations or the experimental data alone. This work establishes a new and practical framework for incorporating general protein dynamics information from simulations of mutants to guide protein engineering.

Toc Graphic

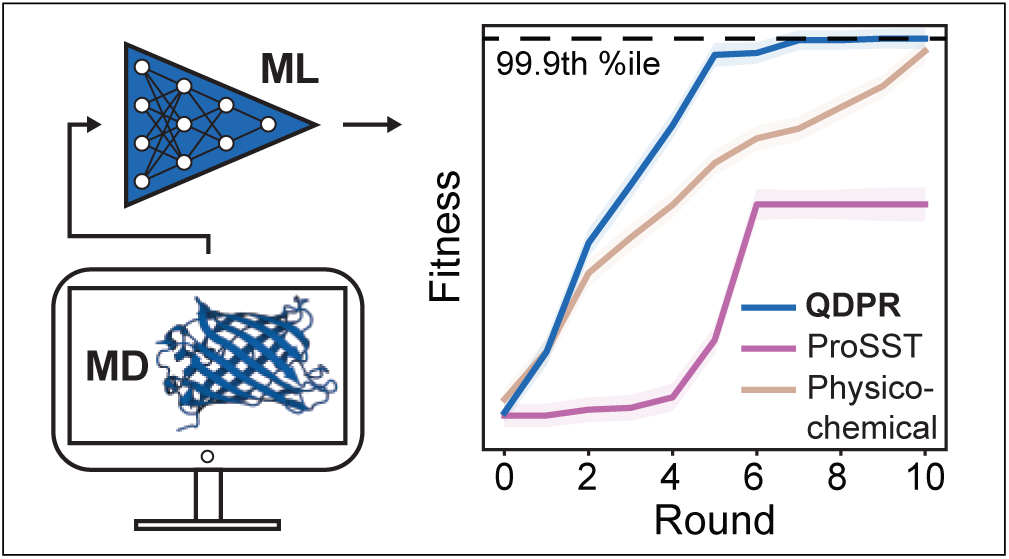

## Introduction and Background

Protein engineering has been greatly accelerated by the application of machine learning (ML) to build mutation effect prediction models: tools trained to accurately predict the effects of one or more mutations on protein functions of interest, such as substrate specificity, thermostability, binding affinity, or any other property.^1^ Despite the promise of this rapidly emerging field, a principal limitation of ML for predicting mutation effects is the availability of appropriate training data. Even highly specialized models making predictions of the effects of low numbers of mutations per protein variant typically require at least hundreds or thousands of high-quality training examples,^2^ which can be prohibitively expensive to obtain. This issue is further compounded for larger numbers of mutations per variant, and models may need to be partially or entirely retrained on new data once more than a handful of mutations have been applied to the initial protein sequence.

Large-scale experimental sequencing approaches like deep mutational scanning have made progress towards meeting the need for large amounts of labeled protein sequence data, but these remain expensive, limited by available transfection and sequencing technologies, and restricted to targets where an appropriate functional assay is available.^3^ Accordingly, a fairly large body of literature has been published in recent years aimed at developing more efficient ML tools for obtaining better variants through fewer experiments. Many strategies have been proposed, but one major commonality is the application of additional information to guide the selection of training data, such as through unsupervised clustering of candidate sequences to diversify training data,^4^ pre-filtering based on predictions of structural stability,^5,6^ or through the use of protein language models or encodings trained on large numbers of unlabeled protein sequences. ^7–10^ Despite the improved performance of all of these approaches over unguided directed evolution or simple ML models, in practice the number of experiments required to obtain highly optimized variants often remains high, and in particular the accurate prediction of epistatic effects (non-additivity in the effects of multiple mutations) presents a significant challenge.^11,12^

Additionally, existing mutation effect prediction ML models lack the ability to offer mechanistic insight, and operate as a black box. In other words, no matter how accurate a model may be at predicting the effect of a mutation on protein function, its impact is limited by the absence of explanations as to *why* those effects arise. If mutation effect prediction models were capable of reporting molecular-level details about the relationship between sequence and function, this would have a potentially transformative impact on the ability of researchers to understand and engineer protein behavior.

Here, I develop a framework for protein mutation effect prediction that can obtain highly optimized protein variants within only a handful of experimental measurements (on the order of tens), while also providing molecular-level explanations of those effects. This approach is based on combining traditional mutation effect prediction models with dynamic, biophysical information obtained from relatively short high-throughput molecular dynamics simulations of protein variants that roughly capture the effects of mutations on local protein dynamics. This method is an evolution of quantified structure-property relationship (QSPR) modeling that I call quantified dynamics-property relationship (QDPR) modeling. I first explain the rationale and design of the method, and then demonstrate that this approach can produce high-fitness variants with very small experimental budgets across two highly distinct proteins and functions that have been used as common models of epistasis:^13^ the *Streptococcus* protein G B1 domain (GB1) and its affinity for binding human IgG, and *Aequorea victoria* green fluorescent protein (*Av*GFP) fluorescence intensity. Then, I show how predictions of protein dynamics provide a framework for interpreting the experimental data in the context of physically meaningful molecular quantities. Excitingly, the model is able to identify key residues mediating protein-protein binding interactions based on simulations of variants of just one protein, by correlating predictions of molecular-level dynamic effects with benchtop binding affinity measurements. This work establishes a new and powerful approach to protein engineering that is synergistic with existing approaches, opens a new avenue for interpreting the dynamic basis of protein function based on small amounts of mutation effect data, and addresses a major challenge in the field by augmenting ML-guided protein engineering with high-throughput atomistic molecular dynamics simulations of protein mutants that do not need to directly measure the engineered property.

## Methods

### Technical rationale

The general outline for the methodology used in this work is shown in Fig. 1. Typical approaches to guiding protein engineering using neural networks focus on models trained directly to predict the functional label of interest (Fig. 1a). This type of model is incorporated here as well, but the central methodological innovation in this work lies in the extraction of large numbers of biophysical features from unbiased molecular dynamics simulations of randomly selected protein variants (Fig. 1b); the training of neural networks to predict each of these biophysical features based on protein sequences (Fig. 1c); and the subsequent training of a downstream score prediction network that takes the outputs of the previous networks as input features (Fig. 1f), which is used to guide the selection of enhanced variants (Fig. 1e).

**Figure 1:**
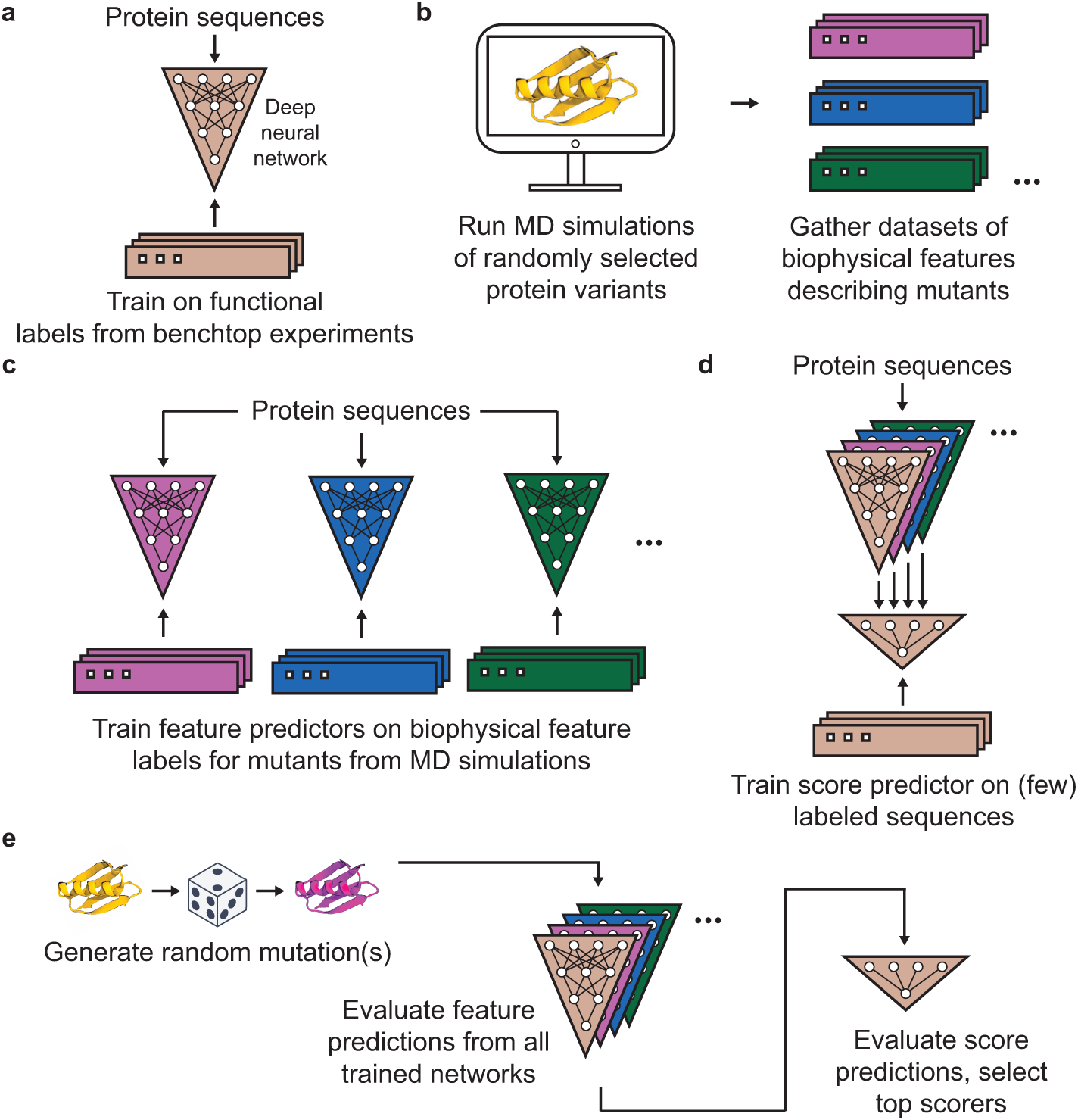
Overview of the method described in this work. **(a)** Training a deep neural network on functional labels describing the property of some protein to be engineered. This represents a typical approach to guiding protein engineering with machine learning (with many recent advances focusing on various learned embeddings for encoding the protein sequences). The three-layer neural network shown in the figure is representative only; in this work, convolutional neural networks were used (see Methods > Training of neural networks). **(b)** Extracting datasets of biophysical features that describe the unbiased simulations of hundreds-to-thousands of randomly selected protein variants. Each color represents a different type of biophysical feature. **(c)** Using those biophysical feature datasets to train a set of additional deep neural networks to make predictions of each feature from protein sequences. **(d)** Training a small score prediction neural network that takes each of the feature predictions as input. The score predictor is trained to predict functional labels from benchtop experiments, just like the model in **(a)**, and uses that model as an input along with the feature predictions. **(e)** Generating random mutations and then evaluating them using each of the trained neural networks. Outputs from the score predictor corresponding to each mutated sequence are used to score sequences. Evaluating the network predictions is fast, so large numbers of possible variants can be efficiently screened without running additional simulations for each evaluated sequence.

Importantly, the simulations themselves are *not* constructed so as to attempt to produce any direct measurement or prediction of the protein property to be engineered. For example, when engineering a protein for improved binding affinity for a particular partner, the binding partner was not included in the simulations at all. Instead, the purpose of the simulations is to collect data describing the molecular biophysical properties of the protein itself, which are then indirectly related to experimental measurements of the target property. The hypothesis guiding this design is that improved protein variants can be selected by biasing the selection process towards variants whose dynamic biophysical properties are suited to achieving desired changes in the target property. This approach also minimizes the amount of information that needs to be known beforehand about the molecular basis of the desired protein property; e.g., it is not necessary to know where or how a partner binds to the target protein in order to apply this method to optimize binding affinity.

### Molecular dynamics simulations

Molecular dynamics simulations were performed using Amber 22.^14^ The GB1 model was based on PDB ID: 1PGA^15^ and the *Av*GFP model was based on PDB ID: 2WUR.^16^ Proteins were modeled in the wild type using the ff19SB force field^17^ and an octagonal box of OPC3 water molecules^18^ with a minimum distance between the protein and the edge of the box of 8 Å. The H++ webserver^19^ was used to determine initial protonation states of titratable residues. Models were minimized over 5000 steps and then heated at constant volume from 100 to 303.15 K over 11,000 2-fs steps. Hydrogen mass repartitioning^20^ with a factor of 3 was applied to enable a longer simulation time step, and the models were equilibrated at constant pressure over 250,000 4-fs time steps. Based on these initial equilibrated models, mutated models were generated using PyRosetta^21^ to apply mutations and PROPKA3^22^ to assess changes in protonation states before being re-minimized and heated using the same workflow as for the wild type model. Mutations were randomly selected (uniform probability across all 19 non-wild type amino acids at each mutated position), with uniform probability of mutating each position across the whole protein. The number of mutations per simulation was also selected uniformly randomly, with either one or two mutations for GB1 or anywhere between one and seven mutations for *Av*GFP (the residues making up the chromophore in GFP were not subject to mutation).

Production simulations for each variant were run for 100 ns (arbitrarily, in a single, continuous trajectory for GB1 or as two, independent 50-ns trajectories for *Av*GFP). One frame was captured for analysis every 10 ps for both proteins. These simulation lengths are fairly short for measuring protein-wide phenomena, and were selected so as to prioritize spending computational resources on a large number of variants while only roughly sampling the effects of each mutations on residue-level dynamics across the proteins, with no expectation of achieving fully convergent sampling.

### Extraction of biophysical features from simulation

Simulations were analyzed using pytraj,^23^ MDTraj,^24^ and MDAnalysis.^25,26^ Before analysis, the first 10% of each trajectory was discarded to allow for equilibration. Five different types of biophysical features were extracted from each GB1 simulation: by-residue root-mean-square fluctuation (RMSF), by-residue Kabsch-Sander backbone hydrogen bonding energy,^27^ by-residue Wernet-Nilsson hydrogen bonding energies,^28^ by-residue Shrake-Rupley solvent accessible surface areas using a probe of radius 1.4 Å,^29^ and the relative weights of the projections of each mutant trajectory onto each of the first 70 components of a PCA decomposition of the motion of alpha carbons from an arbitrarily selected representative trajectory for that protein (70 was chosen semi-arbitrarily as that was the number that captured roughly 95% of the variance in the representative trajectory for GB1; the same number captured roughly 87% of the variance in the representative *Av*GFP trajectory). For the *Av*GFP simulations by-residue global allosteric communication scores^30^ were included as features and Shrake-Rupley terms were excluded. These features – totaling 294 labels for each simulated variant of GB1 and 848 for *Av*GFP – served as training data for the neural networks in the next step.

### Training of feature prediction neural networks

Convolutional neural networks (CNNs) for each protein were trained to predict each of the biophysical features based on the methodology, network architectures, and hyperparameters described for the corresponding sequence CNNs for each protein by Gelman et al. ^2^ As in that work, protein sequences were encoded using a combined one-hot and physicochemical properties encoding based on the amino acid index database, AAindex1.^31^ Each biophysical feature network was trained on data from the same set of 2000 (for GB1) or 1500 (for *Av*GFP) MD simulations, with 124 or 1000 samples, respectively, reserved for validation. Models were trained until the validation mean squared error stopped decreasing, with a patience of 500 epochs, and restoring the model weights with the lowest validation error at the end. This process was repeated three times per model, with only the lowest-error model against the validation set saved for later use. Label values were linearly rescaled to between 0 and 1 before training. Performance benchmarks for each of the trained models are available as Supplementary file pearson_scores.xlsx.

Neural networks trained to directly predict the experimental protein property labels (which we call here the “feature-level” property predictor, to be used as an input to the score predictor; not the score predictor itself, which is described below) were trained in the same manner, except with a variable validation set size reflecting 10% of the total available data and a patience equal to the total number of steps for which the corresponding model was trained in Gelman et al. ^2^ Overall better performance was observed with this approach compared to using no validation set with a fixed number of training epochs.

### Prediction of protein property scores

An additional small neural network was trained to predict the final protein property scores, taking the predictions from the feature prediction networks described above (including the feature-level property predictor) as inputs. This is schematized in Fig. 1d. The score prediction network consisted of eight densely connected units with the Keras 2.13.1 leaky ReLU activation function followed by a single output unit with linear activation. This model was trained over 400 epochs with a learning rate of 0.0001 with a batch size of 8 on all of the available experimental labels corresponding to the sequences passed into the feature prediction networks. This is the same training data used to train the feature-level property predictor (see Fig. 1a,d). Outputs from the score predictor were then used to score unseen variants as described in the following subsection.

We also trained a network combining QDPR with ProSST 2048, a highly performant transformer-based model that incorporates protein language model and 3D structure encodings.^32^ In this case, the only difference in the construction of the model is that the encodings for the feature-level property predictor model are output logits from ProSST 2048^32^ rather than AAindex1 physicochemical encodings.^31^ The ProSST model itself was not retrained. Results reported for supervised ProSST 2048 alone in this work (as opposed to QDPR + ProSST) are direct outputs from the feature-level property prediction model trained in this way.

### Machine learning-guided protein engineering

Taking advantage of very large published deep mutational scanning datasets from Olson *et al.* for GB1^33^ and from Sarkisyan *et al.* for *Av*GFP,^34^ protein engineering campaigns (successive rounds of variant selection where the previous rounds were used to inform the model for the next round) were simulated by drawing labeled sequences from databases in place of conducting experiments directly. Selections of an arbitrary fixed number of 8 (for GB1) or 16 (for *Av*GFP) sequences were drawn for each round, with a different random selection used for the zeroth round of each campaign. For each protein, four strategies were compared: convolutional neural networks using physicochemical encodings (using the model architectures and hyperparameters from Gelman et al.;^2^ supervised ProSST 2048; ^32^ our new QDPR approach that incorporates the biophysical feature predictions from the neural networks trained on simulation data; and QDPR + ProSST 2048 as described in the previous subsection. For each case, 100 independent campaigns were performed in order to sample a range of possible outcomes. In each round, each method was used to evaluate the full set of available labeled sequences (536,084 for GB1 and 45,205 for *Av*GFP, less however many had already been selected up to that point) and the top scorers were selected for the next round.

## Results and Discussion

### Simulated protein engineering campaigns

The results of the simulated protein engineering campaigns in terms of the average normalized discounted cumulative gain (NDCG) scores on each dataset that were achieved by a given round of selection are shown in Fig. 2. NDCG is a popular metric for assessing the quality of ranking metrics that places high importance on correctly identifying highly desirable items (i.e., protein sequences) from within the dataset, and was computed using code from ProteinGym.^38^ QDPR + ProSST was the best overall method by the end of each set of campaigns, although QDPR alone slightly outperformed the combination for GB1 with small amounts of training data. A similar plot for the Spearman rank correlation coefficient, is available as Supplementary Fig. 1.

**Figure 2:**
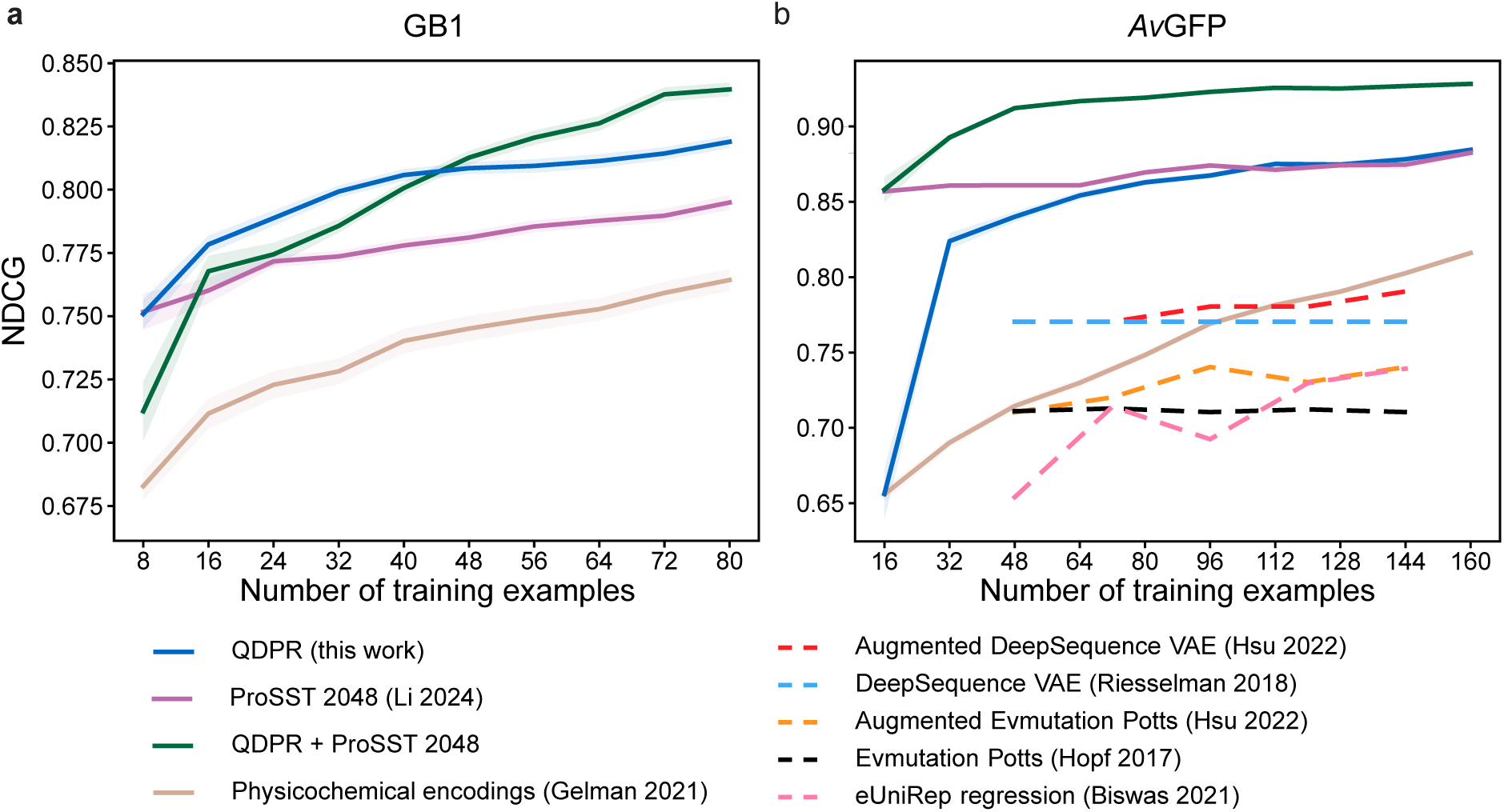
Normalized discounted cumulative gain. Average NDCG scores for each method across the entire dataset based on training data from successive steps of simulated engineering campaigns over 100 independent campaigns each, for **(a)** GB1 and **(b)** *Av* GFP, respectively. Each tick on the horizontal axes accounts for a single additional selection step. Shaded regions represent the standard error of the mean. The sequence CNN with physicochemical encodings approach from Gelman *et al.* 2021^2^ is tan; supervised ProSST 2048^32^ is magenta; QDPR is blue; and QDPR combined with ProSST 2048 is green. Also provided in dashed lines on the *Av* GFP plot for comparison are data from others’ work on a subset of the same dataset, using random sampling instead of simulated campaigns. ^8,35–37^ A version of this figure featuring a comparison on the same dataset and sampling method is available as Supplementary Fig. 2, with similar results.

Because in practice the desired outcome of a protein engineering campaign is usually a single optimized variant, I also looked at the distribution of the highest fitness scores (i.e., binding affinity for GB1 or fluorescence intensity for *Av* GFP) sampled within a given number of rounds with each method. This is a different task than NDCG scores measure because selection of highly optimized variants only requires the identification of positive outliers, regardless of how well other variants are scored. The median fitness scores with standard error are shown in Fig. 3. In both cases, QDPR outperformed alternatives. Notably, despite its relatively stronger performance in NDCG scores compared to physicochemical encodings, ProSST underperformed that method on this metric, indicating that ProSST identifies fewer positive outliers under the low-training data conditions tested. This is also illustrated by the significant negative impact that including ProSST alongside QDPR had on the selection of desirable GB1 variants, and the only very slight impact on selection for *Av* GFP despite large improvements in NDCG.

**Figure 3:**
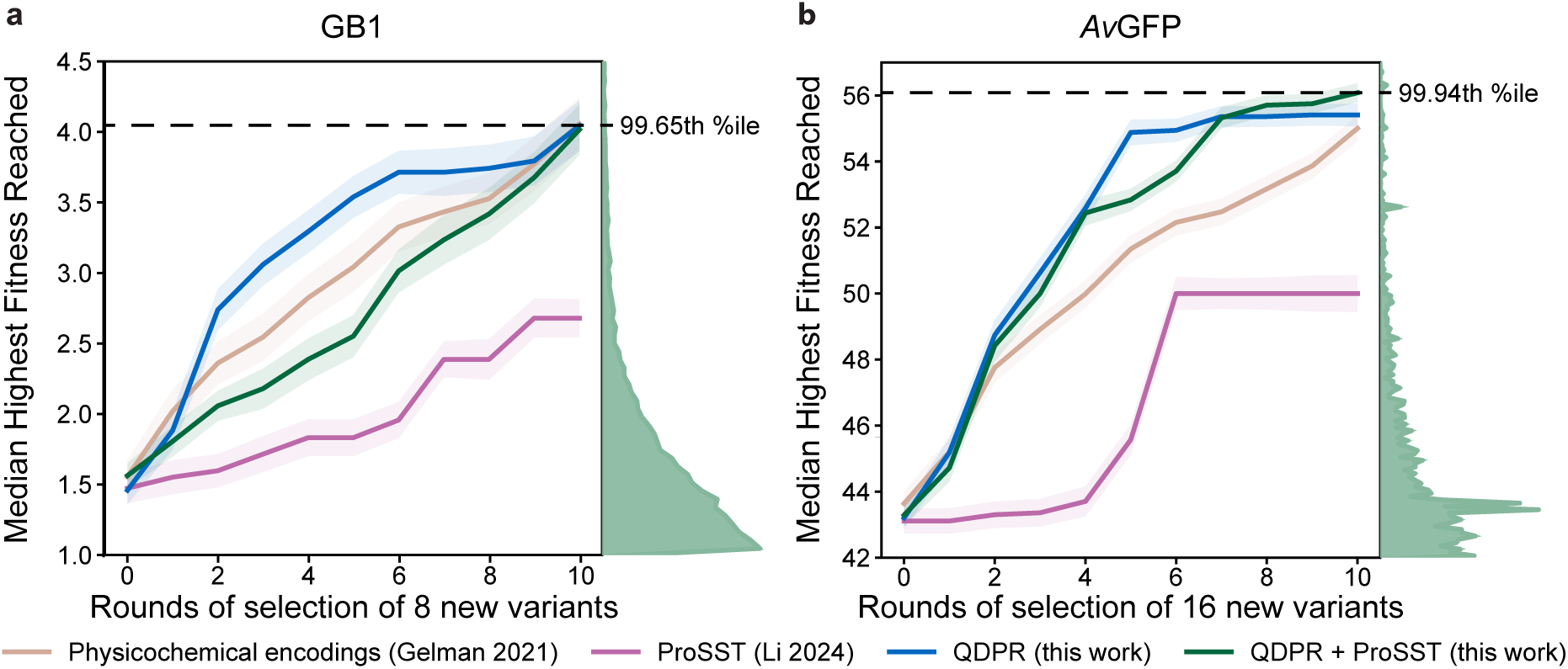
Median highest fitness reached. Each line represents the median highest fitness score reached by a given number of rounds of selection over 100 independent campaigns each, for **(a)** GB1 and **(b)** *Av* GFP, respectively. Shaded regions represent the standard error of the mean. The physicochemical encodings approach from Gelman *et al.* 2021^2^ is tan; supervised ProSST 2048^32^ is magenta; QDPR is blue; and QDPR combined with ProSST 2048 is green. Histograms at the right of each plot represent the distributions of labels in each dataset over the same range of values covered by the plot. The percentile within each dataset of the highest median value reached is indicated with a dashed line.

The *Av* GFP dataset from Sarkisyan *et al.*^34^ is especially valuable for characterizing the ability of models to capture epistatic effects, as it includes a large proportion of variants with many mutations. To characterize the ability of QDPR to model epistatic effects, I assessed the NDCG score as a function of train set size on subsets of the dataset containing one, two, three, four, or five or more mutations, shown in Fig. 4. Across the spectrum of numbers of mutations, QDPR combined with ProSST 2048 remains the clear best performer. Additional data about the sampling of sequences with fluorescence above the WT across different numbers of mutations per sequence are available as Supplementary Fig. 3. QDPR is observed to improve the sampling of desirable sequences across the entire spectrum of numbers of mutations per sequence relative to alternatives, even up to 12-13 mutations.

**Figure 4:**
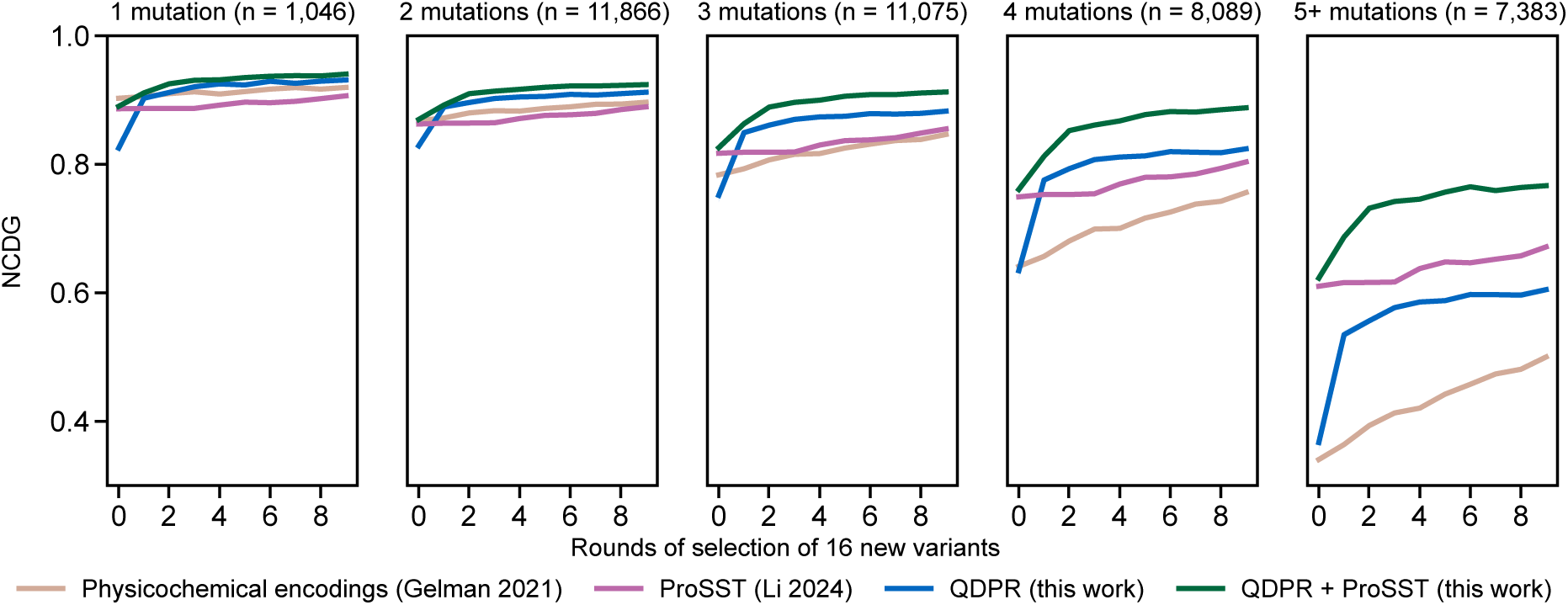
*Av* GFP epistasis. Average normalized cumulative discounted gain for various methods scored on subsets of the *Av* GFP dataset with given numbers of mutations per sequence, with the size of each subset of the dataset indicated. Standard errors of the mean are thinner than the width of the lines. The physicochemical encodings approach from Gelman *et al.* 2021^2^ is tan; supervised ProSST 2048^32^ is magenta; QDPR is blue; and QDPR combined with ProSST 2048 is green.

### Quantified feature importance

Besides improved variant selection, another key advantage of QDPR is that the observed correlations between each biophysical feature and experimentally determined target labels can be used to infer the relative importance of each biophysical feature in the molecular basis of the target protein function. For example, in the case of GB1 binding to human IgG, the bound structure is known (see PDB ID: 1FCC^39^), but IgG was not present in the simulations, providing a test case where the predicted importance of each biophysical feature can be evaluated. As a simple approach to testing this, the “importance” of each of the by-residue features from the MD simulations for a given QDPR campaign after just the first round of GB1 variant selection was quantified as the correlation coefficient between the predicted feature values and the deep mutational scanning property labels for each of the selected sequences up to that point in the campaign. These importances were converted to rank order, and then an overall relative importance score was assigned to each residue by computing the average rank across the features describing that residue. This procedure is schematized in Fig. 5. To visualize the distribution of residue importance scores across the 100 QDPR campaigns, I performed a consensus ranking across each of the QDPR campaigns to obtain the most likely outcome from a single experiment, visualized in Fig. 6. More details of how this consensus ranking was performed are reported in the Supplementary Information. Excitingly, despite this very simple approach the residues with the highest importance ranks (Glu-27 and Lys-31) turn out to be the ones most directly responsible for IgG binding, meaning that even if the GB1 residues that associate with IgG had not been known in advance, they could have been accurately predicted using this approach. Other high-importance residues are hydrophobic residues with their side chains packed inside the protein, with clear roles to play in fold stability. Besides structural evidence, the importance of the identified residues for binding is reflected in the large effect associated with mutating them: whereas the average relative log binding affinity score across all variants in the GB1 dataset is −2.42 ± 2.28 (n = 536,084), the average for all variants wherein either Glu-27 or Lys-31 is mutated is −5.47 ± 0.914 (n = 38,625). Furthermore, the importance of these residues was frequently correctly identified by correlation of biophysical features with just 16 experimentally labeled sequences, even in campaigns where none of those 16 sequences included mutations in either position (82 out of 100 campaigns). Instead, changes in the RMSF, hydrogen bonding energies, and/or solvent-accessible surface areas of these residues were observed in most cases only as consequences of mutations elsewhere in the protein during MD simulations, highlighting the importance of features trained on MD simulations in making accurate predictions with small amounts of experimental data. A consensus ranking that excludes the 18 out of 100 campaigns where mutations to these residues were observed in the training data places Glu-27 and Lys-31 at importance ranks four and three, respectively, behind only hydrophobic residues in the protein core. A similar visualization for *Av* GFP is available as Supplementary Fig. 4, and median importance ranks for each residue across campaigns for GB1 are plotted in Supplementary Fig. 5.

**Figure 5:**
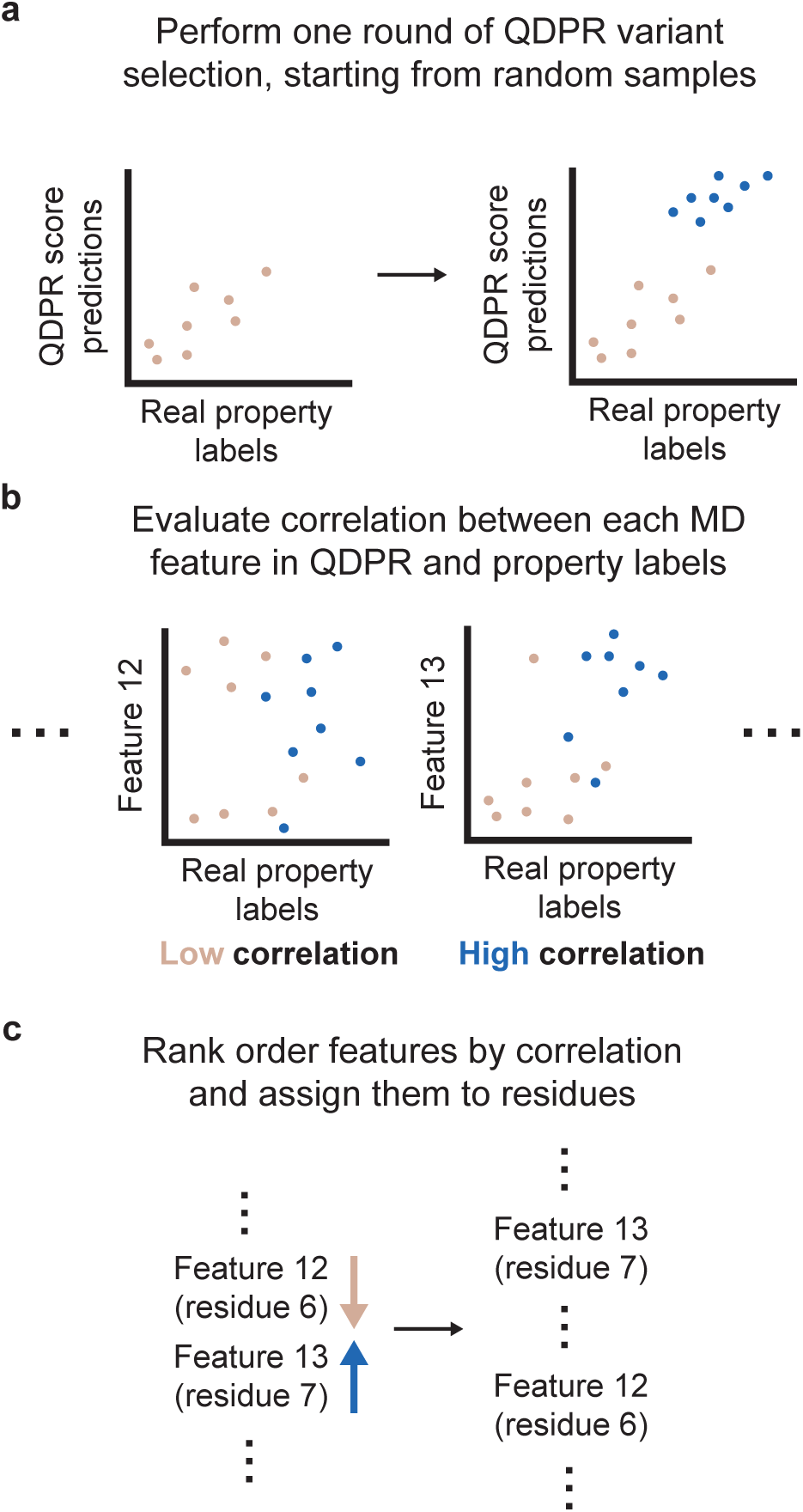
Method for computing residue importance ranks. A schematic describing how importance ranks for features were computed in this work. **a** A single round of variant selection is performed using QDPR. **b** The QDPR prediction is separated into each of its constituent features from MD simulations (here, for example, features “12” and “13”). For each feature, a correlation between the feature predictions and the real property labels is computed. **c** The features are sorted into rank order based on their correlations. Each feature is assigned to a residue (e.g., the RMSF of residue 6 is assigned to residue 6); features that cannot be assigned to an individual residue are discarded. Finally, the ranks for each feature corresponding to a given residue are averaged, and this is interpreted as the overall importance score for that residue.

**Figure 6:**
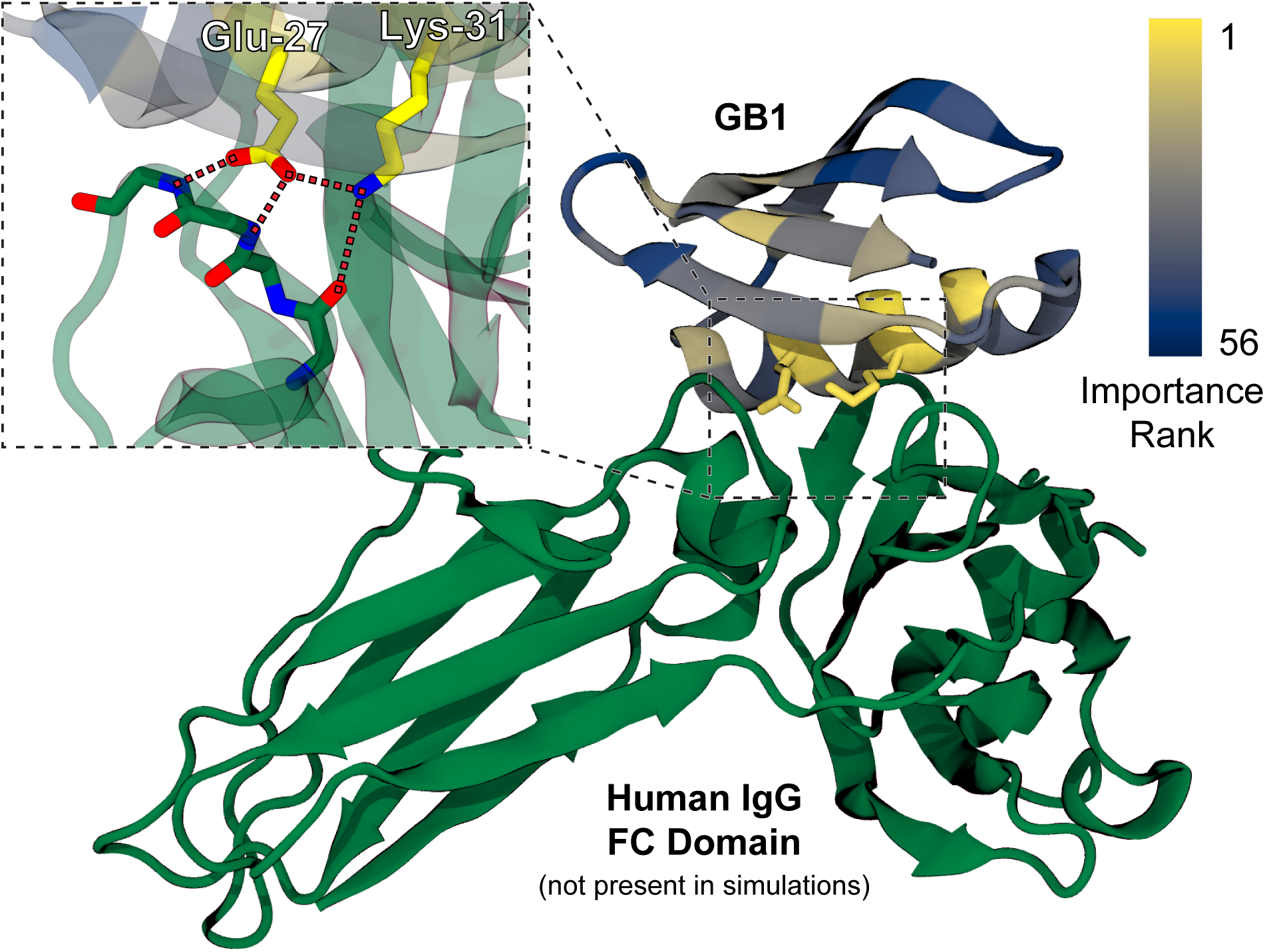
Visualization of residue importance ranks. A portion of crystal structure PDB ID: 1FCC^39^ depicting binding between GB1 the fragment crystallizable (FC) domain of human IgG in green. GB1 is colored by-residue based on the relative importance rank of each position’s by-residue features in a consensus ranking among the 100 QDPR 2000-simulation campaigns after a single round of variant selection each. IgG was not included in the simulations, which were of variants of GB1 in water only. Inset, the two highest-ranked residues – Glu-27 and Lys-31 – are shown to form a key hydrogen bonding network that mediates binding between the two proteins.

### Future directions

Although this work has established the potential usefulness of non-specific protein dynamics data from short, high-throughput simulations of mutants for guiding protein engineering, there remain many unanswered questions and opportunities for further improvement of this method. Many of the parameters and decisions made in obtaining these results were somewhat arbitrary, or otherwise chosen for the sake of simplicity. For example, the length and number of the simulations may be adjustable either up or down in order to obtain more accurate biophysical feature prediction models or to reduce the upfront computational cost before model training. Additionally, there is no reason to expect that the short list of types of biophysical features included in this work represents a complete or optimal description of the relevant biophysical information present in the simulations, and more features could certainly be included to further improve the results and expand the range of features available for interpretation of the molecular basis of experimental observations. In particular, mutation effects that can only be resolved on longer timescales than are observable with the simulations performed here are systematically missed by our approach, though could be measured with longer simulation times. Longer timescale effects will probably be of higher importance in predicting mutation effects on some protein properties, such as those that depend on large conformational shifts, compared to the proteins studied here. Because of the large number of parameters and algorithms that may be improved upon, a more thorough exploration of these questions is left for the future, and significant further improvements in the performance of this approach are almost certainly possible. More thorough bench-marking of performance on a wider range of proteins and target properties, and assessment of performance under various conditions (such as for very highly mutated proteins) will be necessary to more firmly establish the value of this method.

One possible extension of this method that bears special mention is the incorporation of Bayesian active learning, both in the selection of variants for simulations and in the guiding of selection of variants during the subsequent engineering campaign. Bayesian active learning is a subset of ML based on quantifying model uncertainty so as to guide the collection of further model training data that will have the maximum marginal positive impact on model performance. Recent work out of the lab of Frances Arnold has demonstrated that incorporating active learning into an ML-guided directed evolution campaign greatly reduces the number of rounds required to obtain optimized results, and also improves the final optimized sequences compared to ML-guided directed evolution without active learning. ^40^ Similarly, active learning could probably be applied directly to QDPR to improve performance without increasing the number of simulations or experiments required.

Hyperparameters and model architectures in this work were taken from work by others,^2^ but had they not been, features from MD may have presented a better opportunity to select desirable hyperparameters and architectures than is usually possible in few-shot tasks. Because selection of optimized model architectures and hyperparameters is best accomplished using large amounts of training data, suboptimal model design parameters are typically expected when data is scarce. However, optimized model designs are frequently highly transferable across different labels for the same protein. ^41^ QDPR offers a rich set of alternative labels extracted from MD simulations to use in model optimization, likely improving data efficiency compared to the use of unoptimized models for any given protein.

### QDPR in the broader context of ML-guided protein engineering methods

Researchers interested in applying the latest ML methods to improve protein engineering efforts now have a very wide array of choices.^38,42^ QDPR features two distinct advantages compared to alternatives conferred by its leveraging of molecular simulations:

1. **Quantified feature importance.** QDPR is unique among available methods in the ability to provide quantified, molecular-level dynamics predictions to explain *why* any given sequence is highly (or poorly) scored. Additionally, any type of molecular descriptor that can be obtained from an MD simulation can be included in this quantification, without requiring any additional data collection.
2. **Dynamics provides a distinct type of knowledge.** Molecular-scale dynamics information has been largely excluded from state-of-the-art protein engineering approaches to date, despite ample evidence that mutation effects often cannot be explained based only on static structures. ^42,43^ QDPR goes far beyond broad-strokes descriptions of conformational ensembles^44–46^ or biophysical descriptions computed from static structures^47^ and instead leverages atomistic simulation data to accurately predict mutation effects on high-resolution dynamic features. This dimension of information about proteins is distinct from the sources of data that have been used to train other methods, such as unlabeled sequences or structures, suggesting that QDPR could be synergistically combined with other methods. This is evidenced here by our result showing improved NDCG scores when combining QDPR with ProSST (which is trained on unlabeled sequences and AlphaFold structures).

Because of these advantages, QDPR synergizes well with existing protein engineering methods and could be deployed alongside (rather than in place of) other approaches. For example, QDPR might be used to to pre-filter the predictions from another method to remove false positives that are missed by other approaches, and/or to enhance the interpretability of the results of successful protein engineering by any method. However, further work remains to be done in optimizing combined approaches for the selection of positive outliers, rather than merely to improve predictions across whole datasets without translating those improvements into meeting actual protein engineering goals.

The principal limitation to the application of this method in practice is the collection of MD simulation data. As reported here, QDPR is trained on thousands of 50-100 ns simulations per protein, which can represent a substantial investment of computational time depending on the size of the protein target and on access to high-performance computing resources, in particular modern GPUs. In practice, this limitation will largely restrict the usefulness of QDPR for the time being to researchers with access to such resources. Recently, Hou *et al.* introduced SeqDance and ESMDance, two protein language models trained on databases of MD simulations and normal mode analysis data.^48^ These models provide alternatives to QDPR that incorporate some knowledge of protein dynamics without requiring additional simulations, although the performance gains in mutation effect prediction compared to previous protein language models reported in that work are generally modest outside of some specific types of tasks. Striking the optimal balance between taking advantage of simulations of diverse sets of simulations, as in those models, and focusing on simulations of mutants of a single protein at a time, as here, will be an important future direction for research.

Very recent contributions aside, MD simulation in protein engineering has traditionally been limited to exploring the mechanisms of protein function in service of rational or semirational engineering. This approach usually requires that a testable mechanistic hypothesis (e.g., a proposed reaction mechanism, or a suspected allosteric network) is already available, since accessible simulation timescales are limited. However, as a structure is the only information about the protein required to perform the necessary simulations for QDPR, methods like this expand the reach of simulation methods in protein engineering to a broad range of protein targets where the exact relationship between molecular dynamics and function is not known, and may not even be hypothesized. Furthermore, ranked feature importance scores will in such cases prove useful in helping researchers to formulate informed hypotheses of the molecular basis of protein function. This is especially salient in light of the recent successes of protein structure prediction models like AlphaFold^49^ and RoseTTAFold^50^ in expanding the range of proteins for which high-quality structure predictions are available.

Lastly, this method should not be construed as being limited to applications in protein engineering; a similar approach could be used to guide the interpretation of mutation effect datasets by unifying experimental and computational data in order to provide molecular-level interpretations for basic science applications. For example, data describing different phenotypic effects of mutations to proteins *in vivo* could be interpreted in the context of biophysical features from simulations in order to help guide study into the molecular origin of disease. This potential application highlights the implications QDPR has not only for protein engineering, but for protein science in general.

## Conclusion

Here is presented a new method for selecting improved protein variants, and for interpreting the molecular-level effects of mutations on arbitrary experimentally determined properties, based on machine learning applied to biophysical features extracted from atomistic molecular dynamics simulations of protein variants. This method has been shown to permit the selection of improved protein variants across two highly distinct proteins and functions using only very small amounts of experimentally labeled protein variant data, consistently outperforming alternatives across several metrics. Furthermore, this method has been shown to be capable of accurately predicting the key residues implicated in protein function, despite the fact that the simulations are highly general and do not require prior knowledge of the molecular basis of the target property. This work expands the reach of molecular simulation for protein engineering and protein science more broadly and addresses an important gap in the field by incorporating data describing changes in atomistic dynamics upon mutation into the task of predicting mutation effects on protein properties.

## Supporting Information Available

The following files are available free of charge.

- Supporting Information: additional data and methodological details
- pearson_scores.xlsx: Pearson’s correlation coefficients for each trained biophysical model on the validation datasets
- importance_rankings.xlsx: complete lists of consensus ranked importance for features in GB1 and *Av*GFP in descending order after the first round of each QDPR campaign

## Data and Software Availability

All MD inputs, Python scripts for MD analysis and training models, and checkpoint files for models trained on MD data are available at: doi.org/10.5281/zenodo.16377496. A Python program that facilitates the process of performing the simulated protein engineering campaigns is available at https://github.com/Burgin-Lab/qdpr.

## Supporting information

Residue Importance Rankings Datasheet

Model Performance Datasheet

## Acknowledgements

This research was supported by research funding provided by the Thayer School of Engineering at Dartmouth College.

## Author Contributions

TEB is the sole author of this work and was responsible for project ideation, data collection and analysis, and manuscript preparation.

## Competing Interests

The author declares no competing interests.

## Notes

### Competing Interest Statement

The authors have declared no competing interest.

### Summary of Updates

Additional details about the methodology and about the comparison to previously published experimental data have been added. Clarifications to the claims made in the manuscript have been added and additional perspectives based on previously published research are now included.

