## Supplemental Information for "Quantified Dynamics-Property Relationships: Data-Efficient Protein Engineering with Machine Learning of Protein Dynamics"

T. Emme Burgin*

**Thayer School of Engineering, Dartmouth College, Hanover, New Hampshire 03755, United States*

### Spearman correlation coefficients

In addition to the plots of average NDCG scores as a function of training set size, here we provide the same plots for the Spearman rank correlation coefficient as Supplementary Fig. 1. Unlike NDCG, this metric weights the entire dataset evenly, rather than emphasizing the higher-performing sequences more strongly. Under this metric, physichochemical encodings perform especially poorly compared to methods trained on unlabeled sequences for *Av*GFP, but the combination of QDPR and ProSST 2048 remains the highest-performing method tested across both datasets.

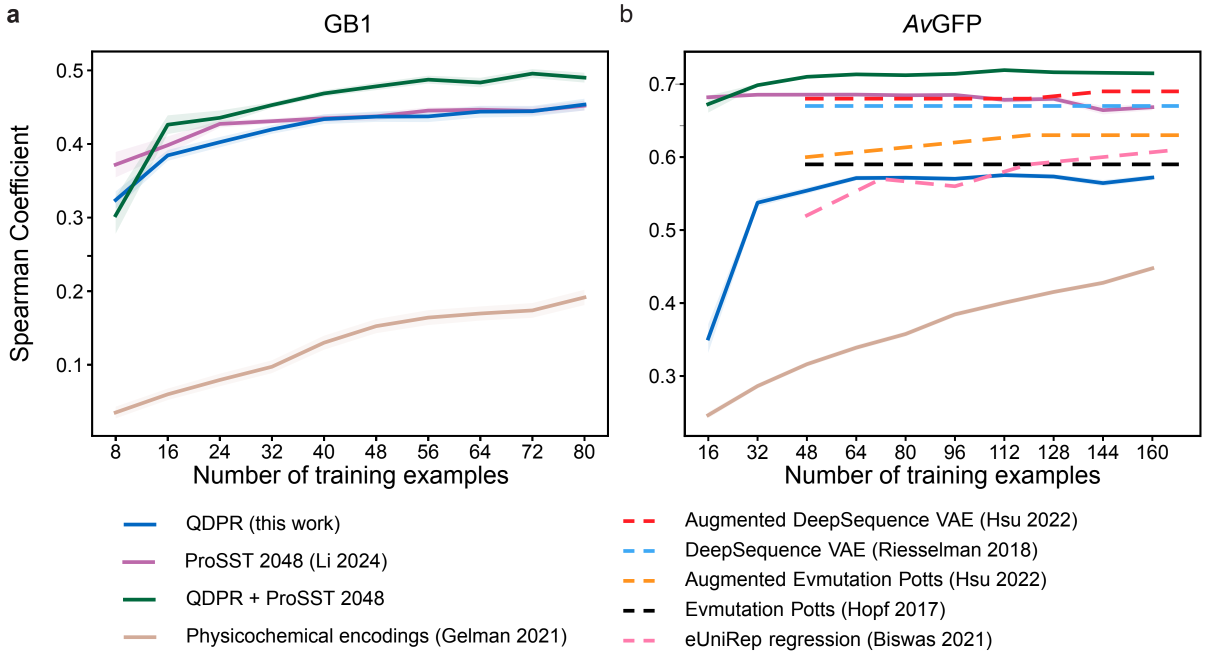


Supplementary Figure 1: **Spearman correlation coefficients.** Average spearman rank correlation coefficients for each method across the entire dataset based on training data from successive steps of simulated engineering campaigns over 100 independent campaigns each, for **(a)** GB1 and **(b)** *Av*GFP, respectively. Each tick on the horizontal axes accounts for a single additional selection step. Shaded regions represent the standard error of the mean. The physicochemical encodings approach from Gelman *et al.* 2021[1] is tan; supervised ProSST 2048[2] is magenta; QDPR is blue; and QDPR combined with ProSST 2048 is green. Also provided in dashed lines on the *Av*GFP plot for comparison are data from others' work on the same dataset that were not collected using simulated campaigns.[3–6]

### Direct comparison of *Av*GFP NDCG scores

Figure 2(b) in the main text includes several normalized discounted cumulative gain scores copied from Hsu 2022;[3–6] however, the method used to select the training data as well as the exact dataset used were different in that work compared to in the present manuscript. Whereas we select training data in successive rounds of 16 new sequences per round, Hsu *et al.* selected new random subsets of their dataset for each number of training examples. Additionally, they used a smaller subset of the same deep mutational scanning dataset, including only mutations in a specific band of residues for which especially strong evolutionary sequence alignment data was available. For the sake of rigorous comparison, we present here a comparison of the average across 100 independent samples each for the four methods that we implemented on that same dataset and with the same sampling method. By comparison to the figure in the main text, the overall trends and ranges of values are extremely similar.

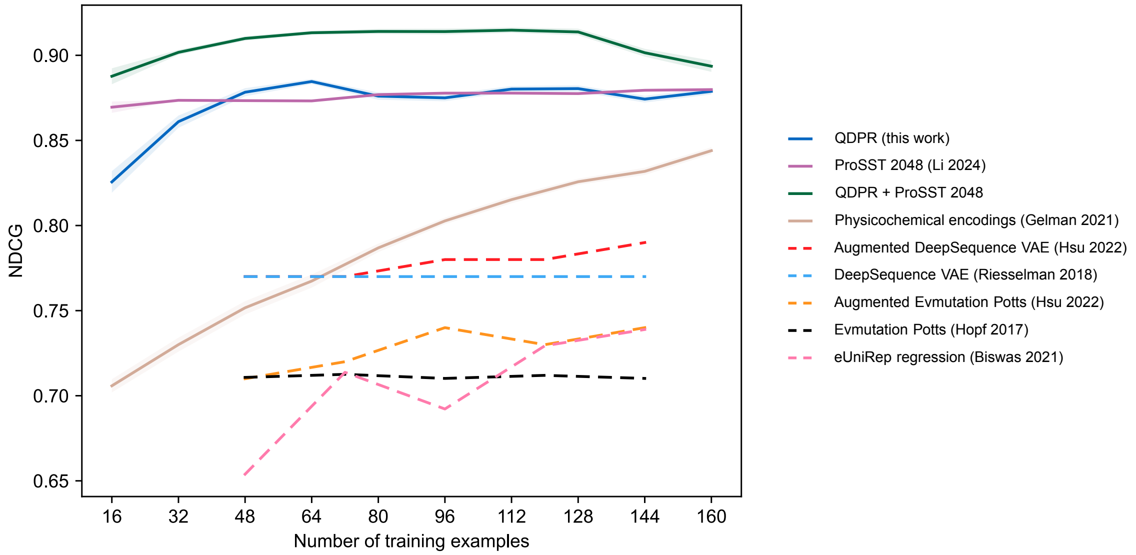


Supplementary Figure 2: **Normalized discounted cumulative gain on Hsu 2022 dataset.** Average NDCG scores on the subsampled *Av*GFP dataset featured in Hsu 2022.[3] Unlike in the main text figures, samples in each round were selected randomly, not by successive rounds of 16 new sequences each. The physicochemical encodings approach from Gelman *et al.* 2021[1] is tan; supervised ProSST 2048[2] is magenta; QDPR is blue; and QDPR combined with ProSST 2048 is green. Also provided in dashed lines on the *Av*GFP plot for comparison are data from others' work on the same dataset that were not collected using simulated campaigns.[3–6]

### Sampling of multiple mutants of *Av*GFP

Figure 4 of the main text concerns the scoring with various methods of subsets of the *Av*GFP dataset containing specific numbers of mutations per sequence, but does not address the actual sampling of desirable highly mutated sequences during simulated protein engineering campaigns. These data are presented here as Supplementary Fig. 3. QDPR is observed to improve the sampling of desirable sequences across the entire spectrum of numbers of mutations per sequence relative to alternatives, even up to 12-13 mutations. ProSST disproportionately prefers less-mutated sequences compared to other methods tested, possibly because it was pre-trained on natural protein sequences and so has learned a preference for more "native-like" sequences, although inclusion of QDPR rescues this bias.

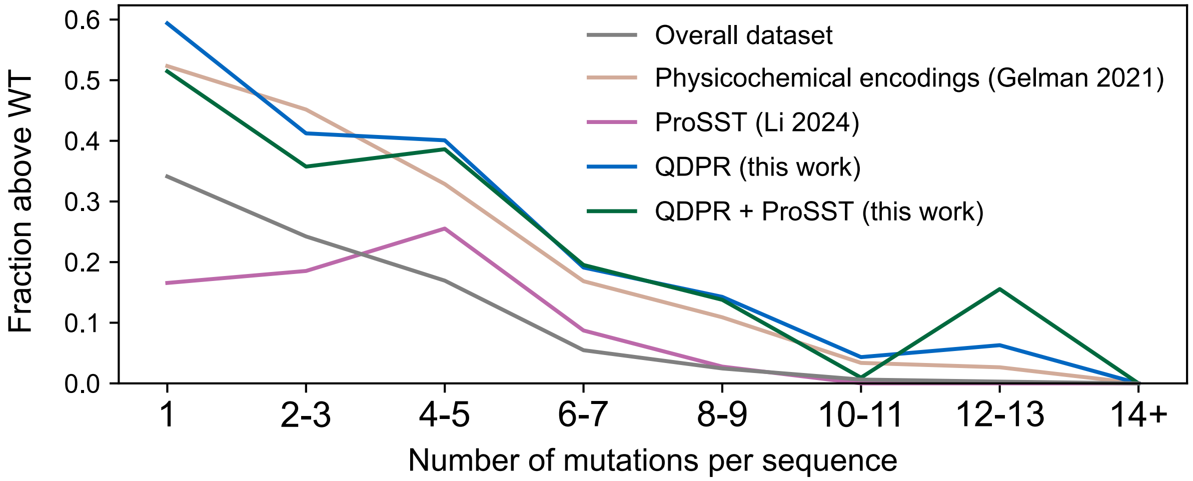


Supplementary Figure 3: **Fraction of sampled *Av*GFP sequences above WT.** Each value represents the fraction of sequences sampled across all 10 steps of the entire set of 100 independent campaigns of protein engineering that are more fluorescent than the wild type *Av*GFP, binned according to the number of mutations per sequence. The fraction of sequences above WT in the overall dataset is shown in gray. The physicochemical encodings approach from Gelman *et al.* 2021[1] is tan; supervised ProSST 2048[2] is magenta; QDPR is blue; and QDPR combined with ProSST 2048 is green.

#### Consensus ranking of feature correlations

More details and results for the consensus ranking process described in the main text are shared here. The consensus ranking was computed using a Python 3 script, the key portions of which are shown here:

```
import itertools
from collections import defaultdict

def calculate_pairwise_preferences(rankings):
    pairwise_counts = defaultdict(int)
    for rank in rankings:
        for i, a in enumerate(rank):
            for b in rank[i+1:]:
                pairwise_counts[(a, b)] += 1
    return pairwise_counts

def calculate_consensus_ranking(pairwise_preferences, items):
    consensus_ranking = list(items)

    consensus_ranking.sort(key=lambda x: sum(pairwise_preferences[(x, y)] -
        pairwise_preferences[(y, x)] for y in items if x != y), reverse=True)

    return consensus_ranking

def pairwise_consensus(file_names):
    rankings = [parse_file(file_name) for file_name in file_names]
    unique_items = set(itertools.chain.from_iterable(rankings))

    pairwise_preferences = calculate_pairwise_preferences(rankings)
    consensus_ranking = calculate_consensus_ranking(pairwise_preferences,
        unique_items)

    return [i for i in reversed(consensus_ranking)]
```

The function *pairwise_consensus* is called on a nested list of feature names in rank order, one for each campaign. The rank order is based on the magnitude of the correlation coefficient between each of the feature values and the corresponding property labels (i.e., GB1 binding affinity or *Av*GFP fluorescence), with more highly correlated features appearing earlier in the ranking. This returns a ranked list of feature names in descending order of importance rank. The complete ranked feature list after the first round of variant selection for each protein is available in the supporting document importance_rankings.xlsx.

A consensus rank visualization for the *Av*GFP campaigns in the main text is shown in Supplementary Figure 4. Across most of the beta barrel structure, residues with side chains facing out (into the solvent) are ranked as less important than their neighbors with side chains facing in (toward the fluorophore). Additionally, residues that are involved in stabilizing the fluorophore – either indirectly by stabilizing the central loop on which it resides, or directly through hydrogen bonding networks, tend to rank quite highly. Other highly important residues may have outsized roles to play in folding.

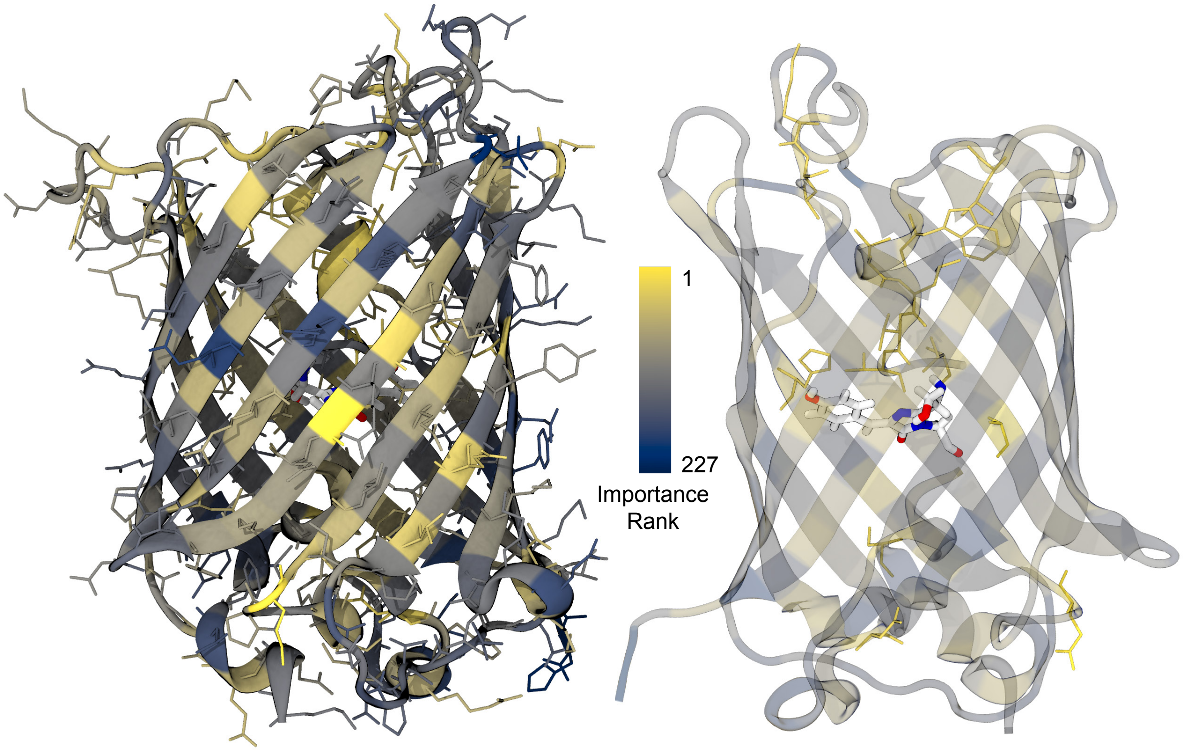


Supplementary Figure 4: **Av*GFP* consensus ranking.** Crystal structure of PDB ID: 2WUR colored according to consensus ranked predicted importance from 100 QDPR campaigns based on results from 1500 simulations after a single round of selection. By-residue importance here is calculated based on Kabasch-Sander backbone H-bond energies, Wernet-Nilsson H-bond energies, and RMSF feature ranks. The chromophore is shown in white and does not have an assigned importance rank. The version at right shows only the top-15 most important residues with ball-and-stick representations.

We also provide the median importance ranks for GB1 residues using the same data that is represented by the consensus ranking in the main text as Supplementary Fig. 5 Higher ranks are considered more important. These data help to characterize the range of values across campaigns.

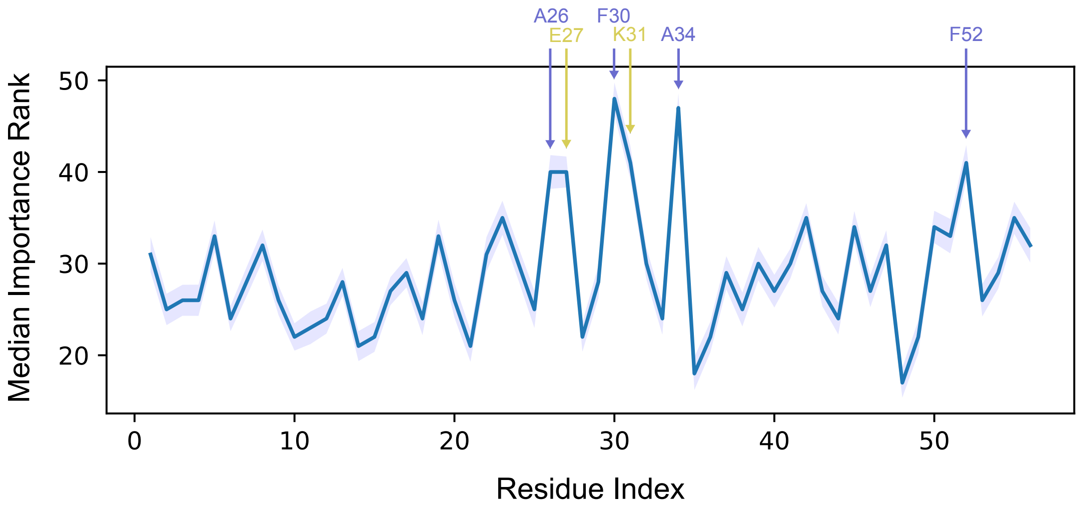


Supplementary Figure 5: **GB1 median importance ranks.** Importance ranks by residue index for QDPR on GB1. Higher ranks are more related to binding affinity. The top six median scores are annotated, and colored according to their role in the protein: periwinkle residues are hydrophobic with side chains inside the protein core, while yellow residues are charged residues at the binding interface. Shaded regions represent standard error.
